# Identification of Actin Filament Interactors in *Giardia lamblia*

**DOI:** 10.1101/2021.05.19.444898

**Authors:** Melissa C. Steele-Ogus, Richard Johnson, Michael MacCoss, Alexander R. Paredez

## Abstract

The deep-branching protozoan parasite *Giardia lamblia* is the causative agent of the intestinal disease giardiasis. Consistent with its proposed evolutionary position, many pathways are minimalistic or divergent, including its actin cytoskeleton. *Giardia* is the only eukaryote known to lack all canonical actin-binding proteins. Previously, our lab identified a number of non-canonical *Giardia lamblia* actin (*Gl*Actin) interactors; however, these proteins appeared to interact only with monomeric or globular actin (G-actin), rather than filamentous actin (F-actin). To identify interactors, we used a chemical crosslinker to preserve native interactions, followed by an anti-*Gl*Actin antibody, Protein A affinity chromatography, and liquid chromatography coupled to mass spectrometry. We found 46 putative actin interactors enriched in the conditions favoring F-actin. Data are available via ProteomeXchange with identifier PXD026067. None of the proteins identified contain known actin-interacting motifs, and many lacked conserved domains. Each potential interactor was then tagged with the fluorescent protein mNeonGreen and visualized in live cells. We categorized the proteins based on their primary localization; localizations included ventral disc, marginal plate, nuclei, flagella, plasma membrane, and internal membranes. One protein from each category was co-localized with *Gl*Actin using immunofluorescence microscopy. We also co-immunoprecipitated one protein from each category and confirmed three interactions. Most of the localization patterns are consistent with previously demonstrated *Gl*Actin functions, but the ventral disc represents a new category of actin interactor localization. These results suggest a role for *Gl*Actin in ventral disc function, which has previously been controversial.

**Importance:** The single-celled eukaryote *Giardia lamblia* is an intestinal parasite that colonizes the small intestine and causes diarrhea and vomiting, which can lead to dehydration and malnutrition. *Giardia* actin (*Gl*Actin) has a conserved role in *Giardia* cells, despite being a highly divergent protein with none of the conserved regulators found in model organisms. Here we identify and localize 46 interactors of polymerized actin. These putative interactors localize to a number of places in the cell, underlining *Gl*Actin’s importance in multiple cellular processes. Surprisingly, eight of these proteins localize to the ventral disc, *Giardia’s* host attachment organelle. Since host attachment is required for infection, proteins involved in this process are an appealing target for new drugs. While treatments for *Giardia* exist, drug resistance is becoming more common, resulting in a need for new treatments. *Giardia* and human systems are highly dissimilar, thus drugs specifically tailored to *Giardia* proteins would be unlikely to have side effects.

## Introduction

Actin is a highly conserved filament-forming protein with essential roles in all eukaryotes that include signaling, motility, membrane trafficking, cell polarity, and cytokinesis (1). Both actin monomers (globular or G-actin) and polymerized actin (filamentous or F-actin) have essential functions in the cell; therefore, the balance between the two forms has a regulatory role in addition to a structural one (2). Eukaryotes throughout the evolutionary tree possess a number of regulators which spatially and temporally control filament formation and depolymerization; their functions include monomer sequestration, filament nucleation or elongation, severing, capping, and crosslinking of filaments (1). Other actin interactors fulfill such roles as linking to organelles and the plasma membrane or moving cargo and generating contractile forces.

The protozoan parasite *Giardia lamblia* is the only eukaryote known to lack all of the canonical actin-binding proteins (3). Due to the lack of conserved interactors that constrain actin evolution, *Giardia* possesses the most divergent actin identified to date, and is only 58% identical to the average eukaryotic actin; in contrast, *S. cerevisiae* actin and human skeletal actin are 87% identical (4, 5). However, *Giardia* actin (*Gl*Actin) retains conserved roles in many cellular processes, including membrane trafficking, cell polarity, and cytokinesis (6, 7). These conserved roles indicate the presence of non-canonical interactors in the proteome (8). The key cellular role of *Gl*Actin as well as its extreme divergence could make it and its interactors potential drug targets. *Giardia* has been designated a neglected disease by the World Health Organization, and giardiasis results in millions of cases of diarrheal disease worldwide each year (9).

Our lab previously identified a number of actin-associated proteins in *Giardia* (10); that study focused on proteins with conserved, identifiable domains whose actin-binding function either may have been overlooked or may not exist in other organisms. Those interactors included microtubule and flagella-related proteins such as p28 and centrin, the chaperone HSP70, the DNA helicase TIP49, the nuclear ARP7, the atypical MAP kinase ERK7, and the regulatory protein 14-3-3.

Much of the *Giardia* genome contains genes annotated as “hypothetical,” as they do not contain any known domains and/or are unique to *Giardia*. Our previous study identified a number of novel actin-binding partners in *Giardia* (10) but did not examine these particular proteins. This study also used a twin-strep tag to affinity purify *Gl*Actin and did not utilize buffer conditions that stabilize filaments; furthermore, the punctate localization of the interactors described were consistent with that of monomeric actin. Therefore, it is likely that this earlier work may have missed proteins that bind exclusively to filamentous actin and is not a comprehensive list of all the actin interactors in *Giardia*.

Here, we used a different approach to discover novel *Gl*Actin interactors for the purpose of identifying those which bind to F-actin. Using a custom anti-*Gl*Actin antibody, Protein A affinity chromatography, and liquid chromatography tandem mass spectrometry (LC -MS/MS), we found a number of novel *Gl*Actin interactors. We categorized a subgroup of these putative interactors based on their localization: marginal plate, flagella, ventral disc, nuclei, membrane, and nonspecific. Notably, no previous *Gl*Actin interactors have been localized to the marginal plate or the ventral disc, suggesting that there may be previously unappreciated roles for *Gl*Actin in these cellular structures.

## Results

### Identification of Non-Canonical *Gl*Actin Interactors

We used a biochemical approach to identify *Gl*Actin-binding proteins, utilizing a chemical crosslinker, anti-*Gl*Actin antibodies, and Protein A beads. For our purification scheme we used buffers developed for canonical actin; whether these buffers are optimal for *Gl*Actin remains untested. Therefore, we also treated the cells with the cleavable chemical crosslinker dithiobis (succinimidyl propionate) (DSP) to ensure the stabilization of native interactions before cell lysis. This molecule reacts with both lysine side chains and the amino terminus of peptides, linking amine groups within 12 angstroms of one another.

In order to differentiate F-actin and G-actin interactors, we lysed wild type cells under these conditions: in G-buffer, expected to favor monomeric actin, followed by DSP treatment, in F-buffer, expected to favor filamentous actin, followed by DSP treatment, or pre-treated with DSP before lysis in F-buffer, expected to stabilize native interactions (Figure 1). We then purified *Gl*Actin using an anti-*Gl*Actin antibody and Protein A affinity chromatography, and subsequently identified the interactors by LC-MS/MS.

**Figure 1:**
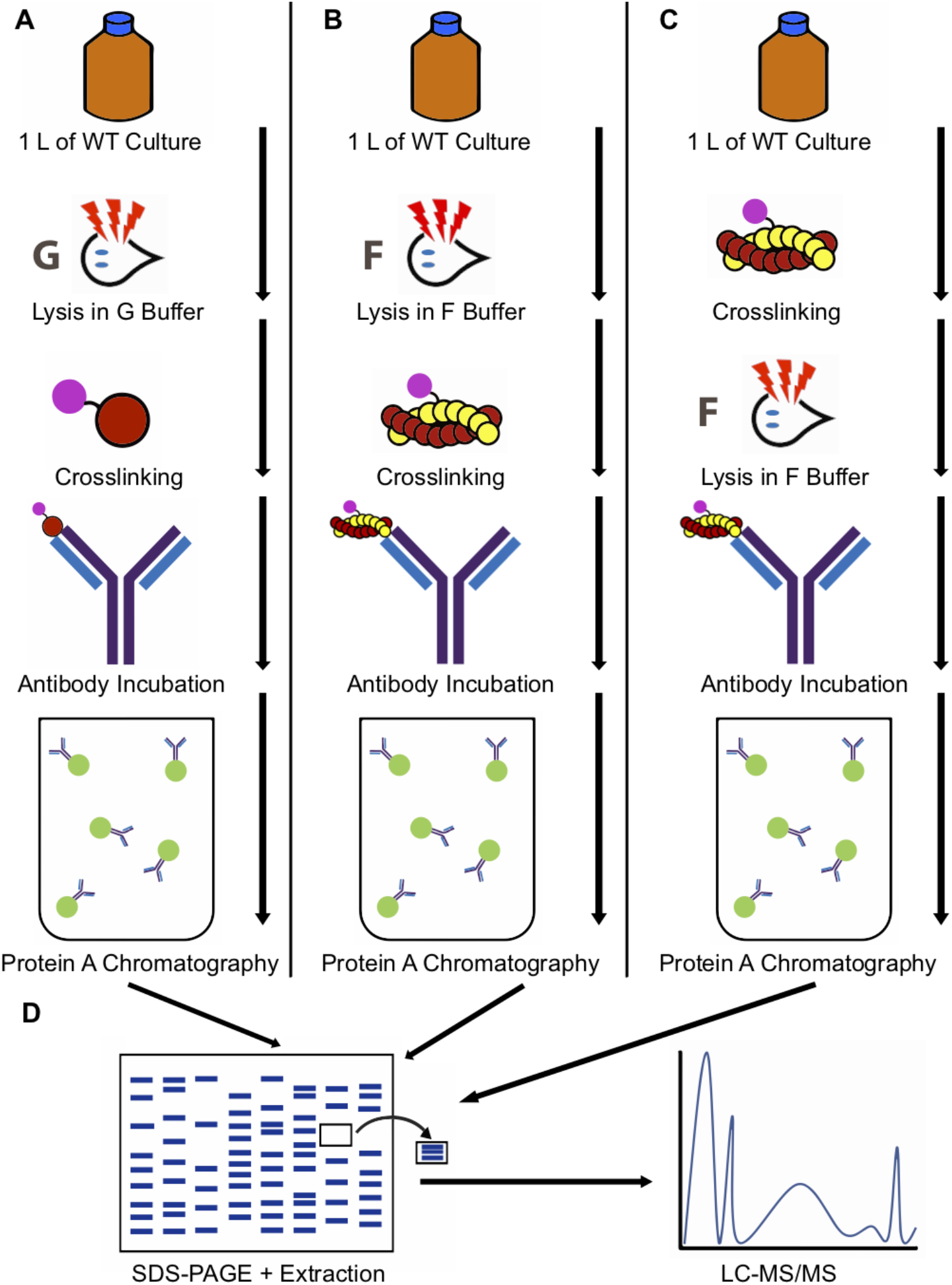
Schematic of methods. 1 L of cells was lysed under one of the following conditions: (A) in G-buffer followed by crosslinking, (B) F-buffer followed by crosslinking, (C) or treated with crosslinker, then lysed in F-buffer. Each lysate was then incubated with an anti-*Gl*Actin antibody followed by Protein A chromatography. (D) The eluate was run on an SDS-PAGE gel, excised, and identified by LC-MS/MS.

Proteins were classified as potential F-actin interactors if they were enriched in either the DSP pre-treated or F-actin buffer conditions in at least two out of three replicates as compared to the G-actin condition. Notably, proteins that had been previously identified as *Gl*Actin interactors, including 14-3-3, p28, GASP-180, and others, were also found in this screen, indicating robustness of our methods (Supplemental Table 1). Proteins enriched in the F-conditions but which are common contaminants, such as metabolic, ribosomal, and proteasomal proteins, were also omitted from our analysis. We also omitted the VPS (variant-specific surface proteins), *Giardia’s* surface glycoproteins which undergo antigenic switching (11, 12). After exclusion criteria were applied, a total of 46 proteins remained (Table 1). Many of these proteins were annotated as “hypothetical proteins,” meaning they lack homologues or known protein domains. Each candidate protein was tagged with the fluorescent protein mNeonGreen and localized in live and/or fixed cells. Five proteins had non-specific localization or low signal (Supplemental Figure 1, Table 1); the others were grouped into the categories discussed below based on their localization (Figure 2). Despite multiple attempts, we were unable to transform GL50803_21423 (beta adaptin) and thus it is absent from our analysis.

**Table 1:** Proteins identified as putative F-Actin interactors

| **ORF Number** | **Identity** | **Localization** |
| --- | --- | --- |
| **Plasma Membrane/Cortical Localization** |  |  |
| GL50803_8560 | Hypothetical | Plasma membrane |
| GL50803_103855 | VPS29 | Plasma membrane |
| GL50803_23833 | VPS35 | Plasma membrane |
| GL50803_4259 | Clathrin Light Chain | Plasma membrane |
| GL50803_8044 | 7-transmembrane domains | Plasma membrane and unidentified compartment |
| GL50803_15591 | Coiled coil protein | Plasma membrane |
| GL50803_14551 | Alpha-6 giardin | Ventrolateral flange and plasma membrane |
| GL50803_17255 | Hypothetical | Plasma membrane, flange, basal bodies |
| GL50803_5810 | Hypothetical | Plasma membrane |
| GL50803_11684 | Leucine Rich Repeat Protein | Plasma membrane |
| **Internal Membrane Localization** |  |  |
| GL50803_9861 | Hypothetical | Diffuse internal membrane |
| GL50803_13930 | Arf3 | Puncta |
| GL50803_22855 | Hypothetical | Vesicles |
| GL50803_12999 | Hypothetical | Puncta and unidentified compartment |
| **Marginal Plate Localization** |  |  |
| GL50803_17153 | Alpha-11 giardin | Marginal plate and axonemal-associated structures |
| GL50803_115478 | Ankyrin repeat, takusan, SMC | Marginal plate and flagellar exit sites |
| GL50803_8854 | Hypothetical | Marginal plate and flagellar exit sites |
| GL50803_4383 | Ankyrin repeat | Marginal plate and axonemal-associated structures |
| GL50803_5800 | Hypothetical | Marginal plate and puncta |
| GL50803_9030 | Ankyrin repeat | Marginal plate and cytoplasmic puncta |
| GL50803_16522 | Hypothetical | Cytoplasmic, marginal disc, nuclei |
| **Nuclear Localization** |  |  |
| GL50803_101212 | Hypothetical | Perinuclear region |
| GL50803_33989 | Hypothetical | Puncta and filaments within nucleus |
| GL50903_14299 | Hypothetical | Diffuse nuclear localization with puncta |
| GL50803_5328 | Sigma adaptin (AP2 complex) | Diffuse nuclear localization and plasma membrane |
| GL50803_102813 | Ankyrin repeat | Nuclear, marginal plate, cytoplasmic |
| GL50803_14213 | Hypothetical | Diffuse nuclear localization with puncta |
| **Flagellar Localization** |  |  |
| GL50803_9848 | Dynein light chain | Internal portions of flagella, basal bodies |
| GL50803_33685 | Armadillo repeat | Flagella |
| GL50803_15097 | Alpha-14 giardin | Flagella |
| GL50803_13774 | Hypothetical | Flagellar exit sites |
| GL50803_4463 | Dynein light chain | Flagella |
| GL50803_15124 | Dynein light chain | Flagella |
| **Disc localization** |  |  |
| GL50803_4239 | Hypothetical | Ventral disc, cytoplasmic |
| GL50803_24451 | FGF | Flagella, basal bodies, lateral crest |
| GL50803_8726 | Hypothetical | Ventral disc, midbody |
| GL50803_27925 | Ankyrin repeat | Ventral disc, except ventral groove |
| GL50803_113622 | Ankyrin repeat | Ventral groove |
| GL50803_16844 | Hypothetical | Ventral disc: ventral groove and overlap zone |
| GL50803_17230 | Gamma-giardin | Ventral disc |
| GL50803_11354 | Hypothetical | Flagella, lateral crest |
| **Non-specific localization** |  |  |
| GL50803_10038 | Alpha-18 giardin | Diffuse non-specific |
| GL50803_10429 | Wos2 protein | Diffuse non-specific |
| GL50803_7323 | Hypothetical | Diffuse non-specific |
| GL50803_6242 | Translationally controlled<br>tumor protein-like | Diffuse non-specific |
| GL50803_21423 | Beta-adaptin | N/A |

**Figure 2:**
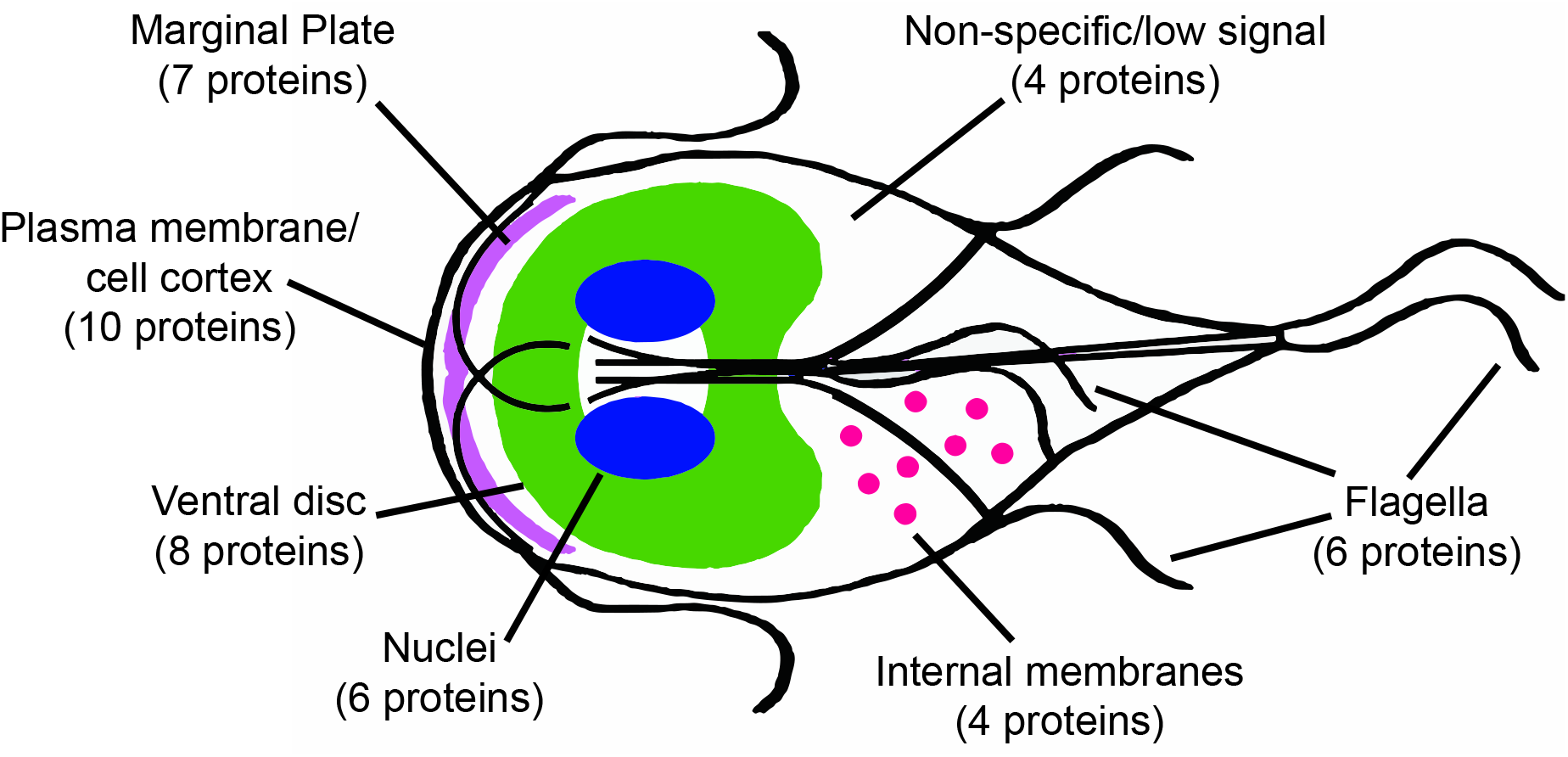
Localizations of *Gl*Actin Filament Interactors. Diagram of *Giardia* cell, with numbers of interactors localizing to each structure in parentheses. Shown are the nuclei (blue), ventral disc (green), marginal plate (purple), flagella and axonemes (black), internal membranes, (pink), plasma membrane (black), and non-specific/low signal. Ventral flagella are shown in cyan for contrast.

### *Gl*Actin Interactors in the Nuclei

*Giardia* has two transcriptionally active nuclei, to which *Gl*Actin localizes (6). Previous studies identified conserved *Gl*Actin interactors in the nuclei and treatments altering the phosphorylation state of *Gl*Actin affected nuclear size (10, 13). In model organisms, actin has multiple roles in the nucleus, including chromatin remodeling and regulating transcription (14).

Our screen discovered six nuclear localized proteins, each with a distinct sublocalization (Figure 3). GL50803_5238 (sigma adaptin, a component of the AP-2 complex) appeared in puncta within the nuclei and in association with the cell cortex. GL50803_102813 (ankyrin repeat protein), a protein upregulated during early encysation (15), was also localized in the cytoplasm, with some large puncta, as well as a slight enrichment in the marginal plate. The protein appears somewhat more concentrated towards the anterior of the nuclei. GL50803_GL14299 (hypothetical) appeared to be associated with the nuclear envelope in puncta. GL50803_33989 (hypothetical) was previously tagged with a C-terminal GFP tag and formed inclusion bodies in cells (16, 17), so we tagged this protein N-terminally under its native promoter. Under these conditions, GL50803_33989 localized to the nuclei, forming structures which may be filamentous. The perinuclear localization of GL50803_101212 (hypothetical) may represent the perinuclear endoplasmic reticulum. In contrast, GL50803_14213 (coiled coil domain) has no puncta or other sub-pattern within the nuclei.

**Figure 3:**
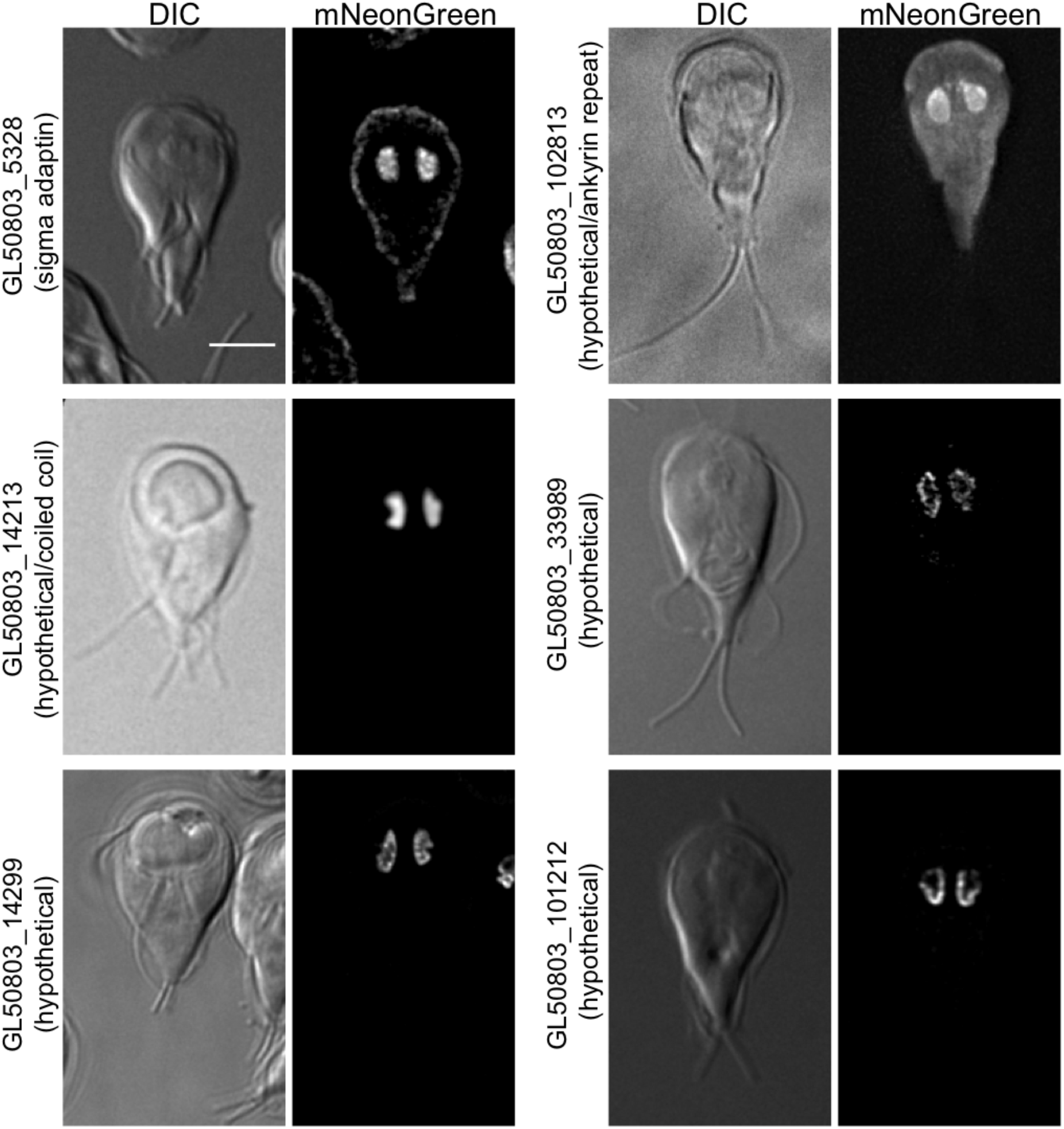
Putative *Gl*Actin interactors with nuclear localization. Six proteins identified localized to *Giardia’s* nuclei based on the localization of mNeonGreen fusions. GL50803_5238 localized to the plasma membrane as well as the nuclei. GL50803_102813 also localized to the marginal plate and cytoplasm, with some puncta. GL50803_GL14299 and GL50803_33989 appeared closer to the nuclear envelope, while GL50803_101212 localized to the perinuclear region. GL50803_14213 was evenly distributed within the nuclei. Scale bar, 5 μm.

### *Gl*Actin Interactors in the Flagella

*Giardia’s* four sets of flagella have different cellular roles and undergo a complex developmental cycle (18, 19). Actin is a key component of flagella; in Chlamydomonas, six of the seven inner dynein arms form a complex with actin (20). Additionally, actin is an important regulator of intraflagella transport and the regulation of flagella length (21). Anti-*Gl*Actin immunostaining showed its clear localization within the flagella, and as a helix that surrounds the caudal flagella, similar to actin organization of the sperm midpiece (22). Our previous study identified six axonemal dynein heavy chains in addition to p28 as *Gl*Actin interactors (10). Thus, finding six flagellar-localized proteins in this screen is consistent with previous knowledge (Figure 4). Three of these candidate *Gl*Actin interactors are dynein light chains; while all appeared in the cytoplasm in low levels, each had their own unique flagellar localization pattern. GL50803_15124 localized to the entirety of the flagella axonemes, while GL50803_9848 appeared primarily associated with the cytoplasmic portions of axonemes. GL50803_4463 had similar axonemal localization, and additionally localized in patches throughout the cell in low levels. GL50803_15097 (alpha-14 giardin) enrichment was less intense in the anterior flagella than the other three flagella pairs. GL50803_33685 (armadillo repeat) appeared throughout the flagella but was particularly enriched in the axonemes. In contrast, GL50803_13774 (hypothetical) appeared only in the flagellar pores, the structures where the flagella exit the cell.

**Figure 4:**
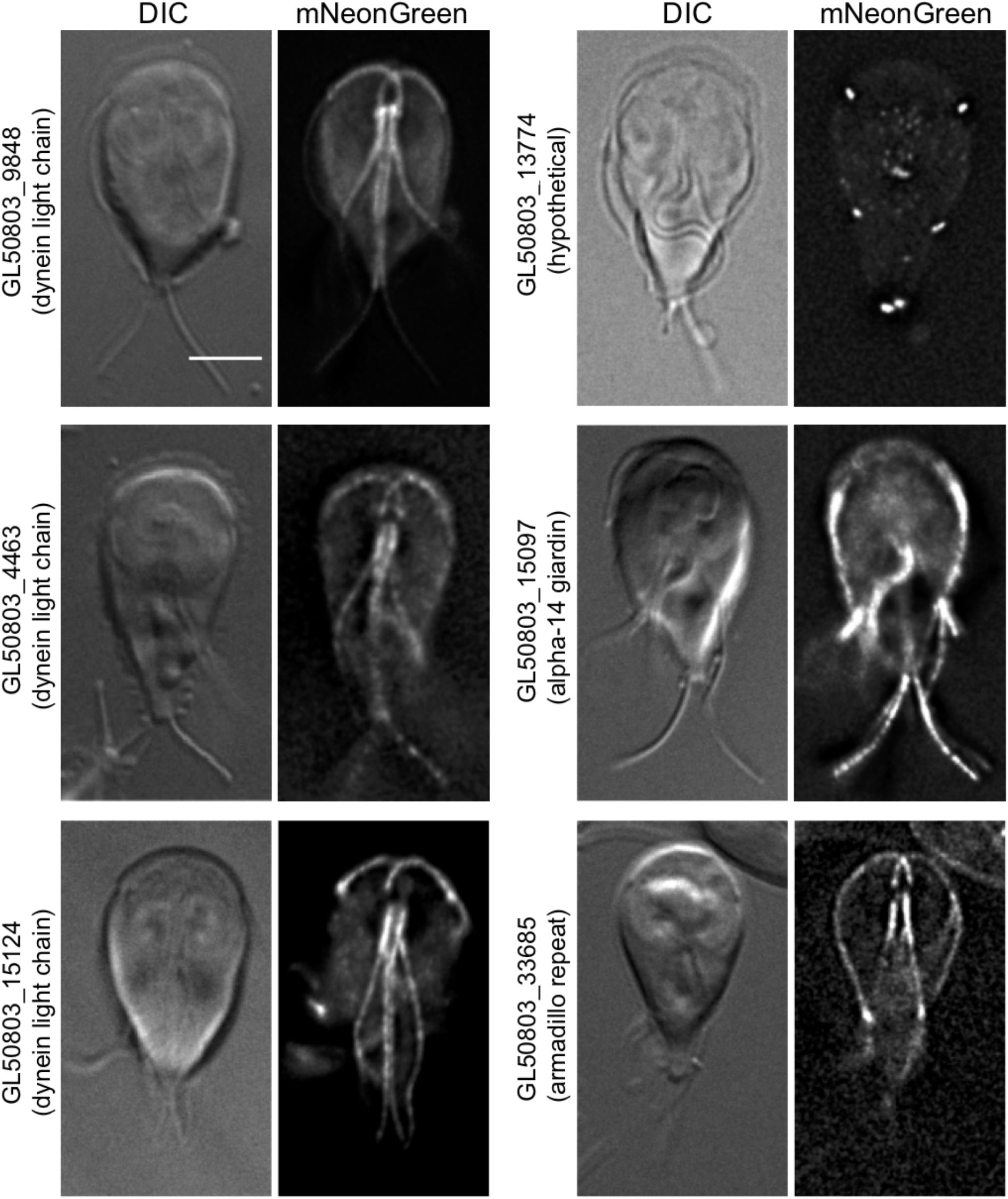
Putative *Gl*Actin interactors with flagella localization. Six *Gl*Actin interactors localized to flagella or flagella-related structures based on the localization of mNeonGreen fusions. GL50803_15124 had a general flagellar localization. GL50803_4463 was enriched in the central axonemes, while GL50803_9848 was enriched in the basal bodies. GL50803_15097 was enriched in posterolateral, ventral, and caudal flagella. GL50803_33685 also appeared slightly in the plasma membrane. GL50803_13774 was only present in the flagellar pores. Scale bar, 5 μm.

### *Gl*Actin Interactors in the Marginal plate

Seven proteins in our screen localized to the marginal plate (Figure 5), a lattice-like structure at the anterior of the cell which is associated with the axonemes of the anterior flagella (23). Ultra-high resolution scanning electron microscopy showed flexible filaments at the plate (24, 25), and immunofluorescence revealed enriched *Gl*Actin there (10, 25). Little else is known about the function or composition of the marginal plate; it has been suggested to have a role in attachment, but this has remained an unexplored area of research (6, 8, 26)

**Figure 5:**
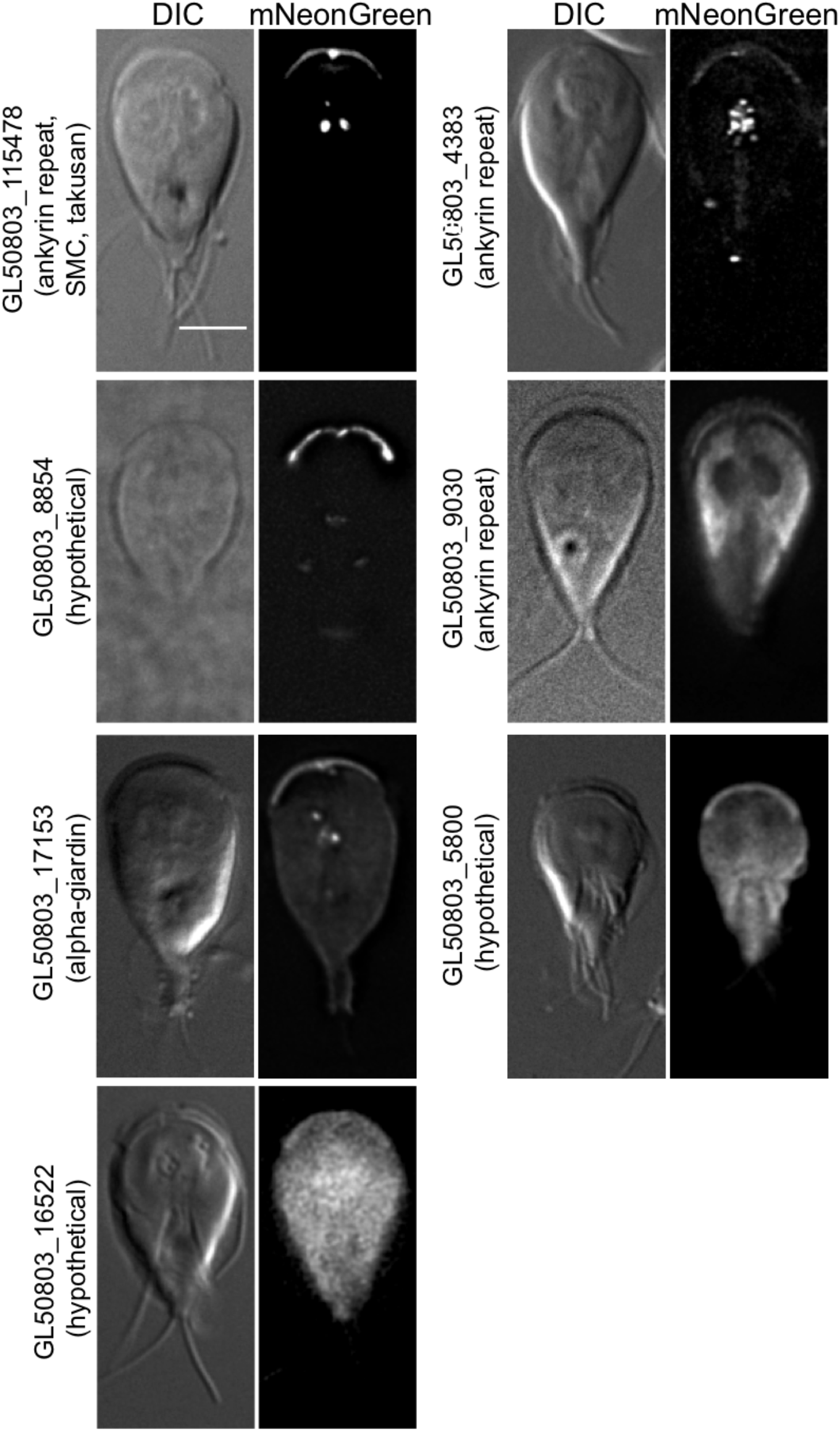
Putative *Gl*Actin interactors with marginal plate localization. Seven *Gl*Actin interactors localized to the marginal plate based on the localization of mNeonGreen fusions. GL50803_115478, GL50803_17153, and GL50803_4383 also appeared in axoneme-associated structures. GL50803_8854 was enriched in the flagellar pores. GL50803_5800 and GL16522 were only slightly enriched in the plate. GL50803_9030 was additionally observed in linear arrays that may be filaments at the anterior ventrolateral flange and also localized throughout the cytoplasm with enrichment in the lateral shields, but was excluded from the nuclei. Scale bar, 5 μm.

Notably, multiple proteins in the marginal plate also localized to microtubule-based structures. For instance, GL50803_8854 (hypothetical) also localized to the flagellar pores, GL50803_17153 (alpha-11 giardin) appeared faintly in the disc (Supplemental Figure 2), and GL50803_5800 (hypothetical) was slightly enriched in the axonemes. GL50803_17153, GL50803_4383 (ankyrin repeat protein), and GL50803_115478 (ankyrin repeat) also appeared as puncta or linear stretches near the axonemes, a localization reminiscent of the mitochondria-like mitosomes (27, 28). These puncta were not present in all cells, and were inconsistent in number (Supplemental Figure 2).

In addition to their marginal plate localization, GL50803_5800, GL50803_16522, and GL50803_9030 (ankyrin repeat) were distributed throughout the cytoplasm. GL50803_16522 also localized to the nuclei. GL50803_9030 also appeared to be more concentrated in the anterior of the cell in the ventrolateral flange and in the lateral shield, the ventral region of the plasma membrane. Both of these structures make contact with the surface of substrates during attachment (29).

### *Gl*Actin Interactors in Internal Membranes

Although *Giardia* lacks many organelles, including a Golgi, lysosomes, and peroxisomes, it still has a defined endomembrane system. *Gl*Actin is important for receptor-mediated endocytosis, membrane remodeling during abscission, and the trafficking of cyst wall protein during encystation (6, 8, 30).

Several of our novel *Gl*Actin interactors have localization consistent with membrane-bound compartments (Figure 6). Intriguingly, GL50803_12999 (hypothetical) localized to a previously undescribed organelle as well as puncta that could be peripheral vacuoles and the ER. GL50803_9861 (hypothetical) also had a punctate localization which could represent small vesicles. Apart from the previously mentioned GL50803_101212, which may be in the perinuclear ER, two proteins localize to internal structures which we suspect to be the endoplasmic reticulum: GL50803_22855 (hypothetical) and GL50803_13930 (Arf3). GL50803_22855 (hypothetical) appeared to be localized throughout the entirety of the organelle, whereas the puncta of GL50803_13930 (Arf3) are consistent with the ER exit sites (31).

**Figure 6:**
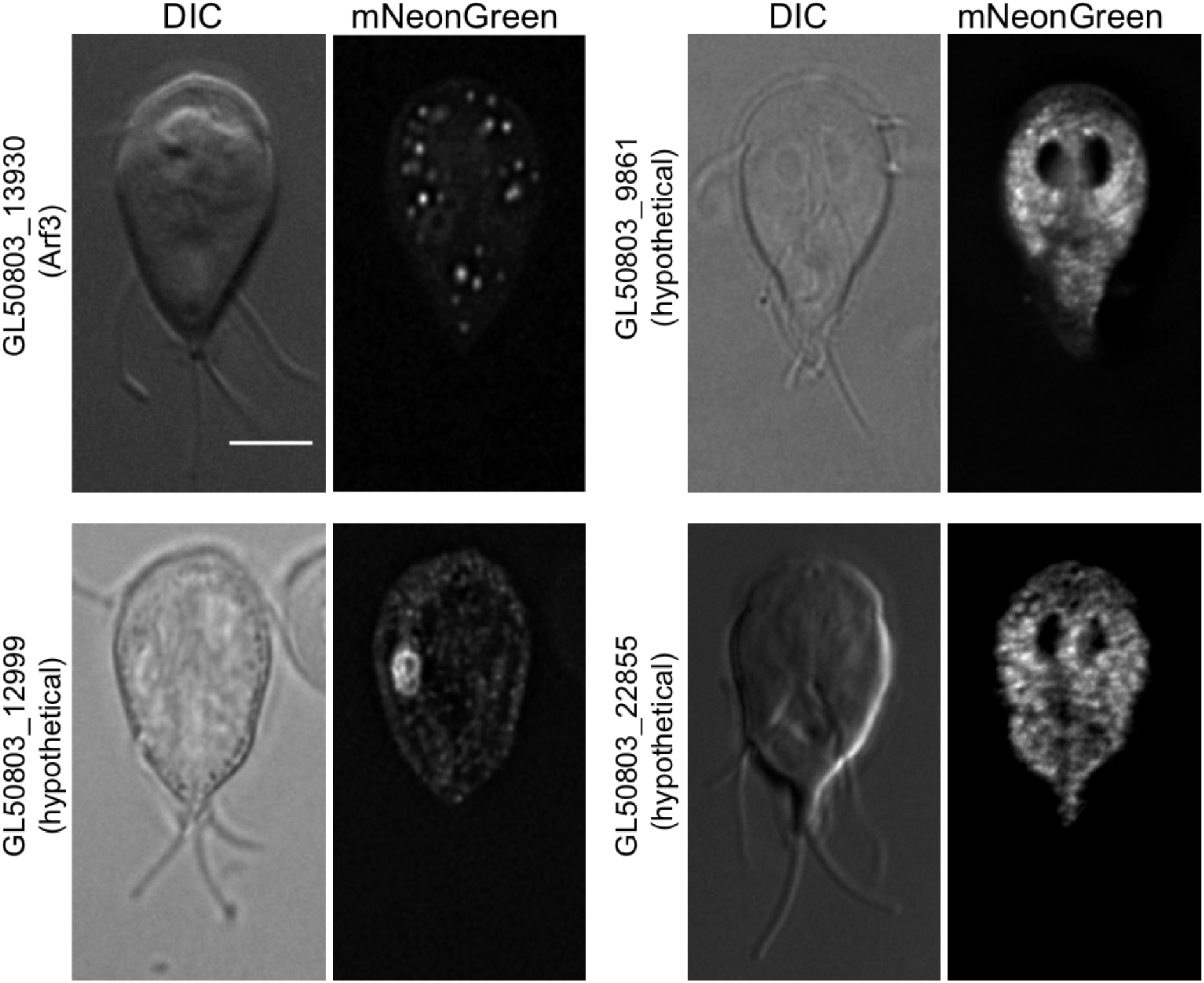
Putative *Gl*Actin interactors in the endomembrane system. Four *Gl*Actin interactors localized to internal structures likely to be the endomembrane system. GL50803_22855 and GL50803_9861 were widely distributed throughout the cytoplasm, GL50803_13930 was concentrated in a few stable puncta, and GL50803_12999 localized to cytoplasmic puncta and a large, oblong organelle. Except for Arf3, these proteins are sometimes seen in linear arrays that could also indicate a connection to actin filaments. Scale bar, 5 μm.

### *Gl*Actin Interactors at the Cortex and Plasma Membrane

We found ten putative interactors which localized to the cell cortex, composing the largest group in the screen. This is consistent with *Gl*Actin and actin in general having roles in regulating cell shape, cell polarity, and membrane trafficking (Figure 7). Despite a shared generalized localization, each putative interactor had its own individual pattern. The internal membrane localization of GL50803_8044 (7 transmembrane domain) localizes to both the plasma membrane and ER. In this case we are confident the protein is in the ER, as transmembrane proteins are necessarily trafficked through this organelle. Both GL50803_14551 (alpha-6 giardin) and GL50803_17255 (hypothetical) appeared in the ventrolateral flange, a thin membranous structure that protrudes from the plasma membrane and contributes to parasite attachment (32, 33). GL50803_23833 (VPS35), GL50803_103855 (VPS29), GL50803_8560 (hypothetical), GL50803_15591 (coil coil protein), GL50803_4259 (clathrin light chain), and GL50803_11684 (leucine rich repeat protein) all had a punctate localization at the cell cortex, likely in association with the plasma membrane.

**Figure 7:**
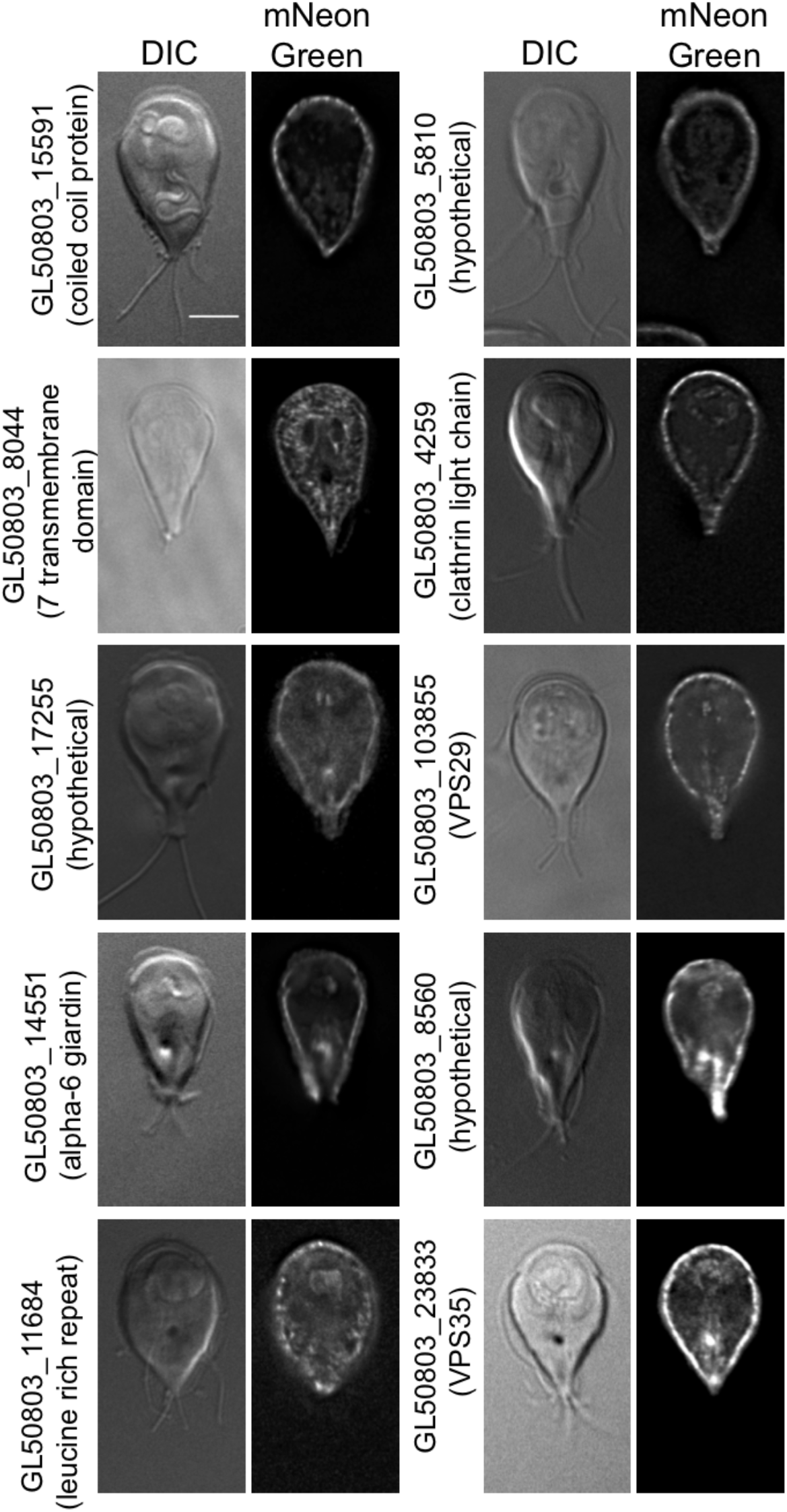
Putative *Gl*Actin interactors with cortical or plasma membrane localization. Ten *Gl*Actin interactors localized to the cell cortex based on the localization of mNeonGreen fusions. Clathrin light chain GL50803_4259 is expected at the cell cortex. GL50803_5810 also had a slight localization to the nuclear envelope and GL50803_8044 localized to the ER. Both GL50803_14551 and GL50803_17255 extended into the ventrolateral flange; the latter was also enriched in the basal bodies. GL50803_103855, GL50803_23833, and GL50803_8560 had a slightly punctate distribution within the plasma membrane. GL50803_15591 and GL50803_11684 were even more punctate, with individual puncta clearly visible. Scale bar, 5 μm.

### *Gl*Actin Interactors in the Ventral Disc

The ventral adhesive disc is a microtubule-based structure unique to *Giardia*, which allows cells to attach to the small intestine of their host, or to the side of a culture tube *in vitro*. The dome-shaped disc is composed of roughly 100 microtubules which form a stable sheet that wraps into a spiral and overlaps itself (34). In the center of the disc is the bare area, a region devoid of microtubules but containing membrane-bound vesicles involved in membrane trafficking (29). *Gl*Actin’s role in the disc, if any, has heretofore been controversial (35).

Eight putative *Gl*Actin interactors localized to the disc, representing the second largest group of putative interactors. GL50803_8726, GL50803_16844, GL50803_4239, and GL50803_17230 had not been localized when we started this work, but have since been identified as disc-associated proteins (DAPs) (17), they are included here for the sake of completeness. GL50803_27925, GL50803_113622, GL50803_24451, and GL50803_11354 are new DAPs.

The localizations within the ventral disc varied widely (Figure 8). Four of these proteins have no recognized domains, including GL50803_8726, which localized to the entirety of the ventral disc in addition to the median body, a reservoir of microtubules supporting rapid disc assembly during mitosis (36). GL50803_16844 (hypothetical), GL50803_4239 (hypothetical), and GL50803_1720 (gamma-giardin) were also localized to the entirety of the disc, but enriched in various regions within it. Both GL50803_11354 (hypothetical) and GL50803_24451 (FGF domain) had less distinct localizations. The remaining two disc-localized proteins have ankyrin-repeat domains, and coincidentally had inverse patterns of localization: while GL50803_27925 appeared everywhere in the disc except the ventral groove, GL50803_113622 localized to only the ventral groove.

**Figure 8:**
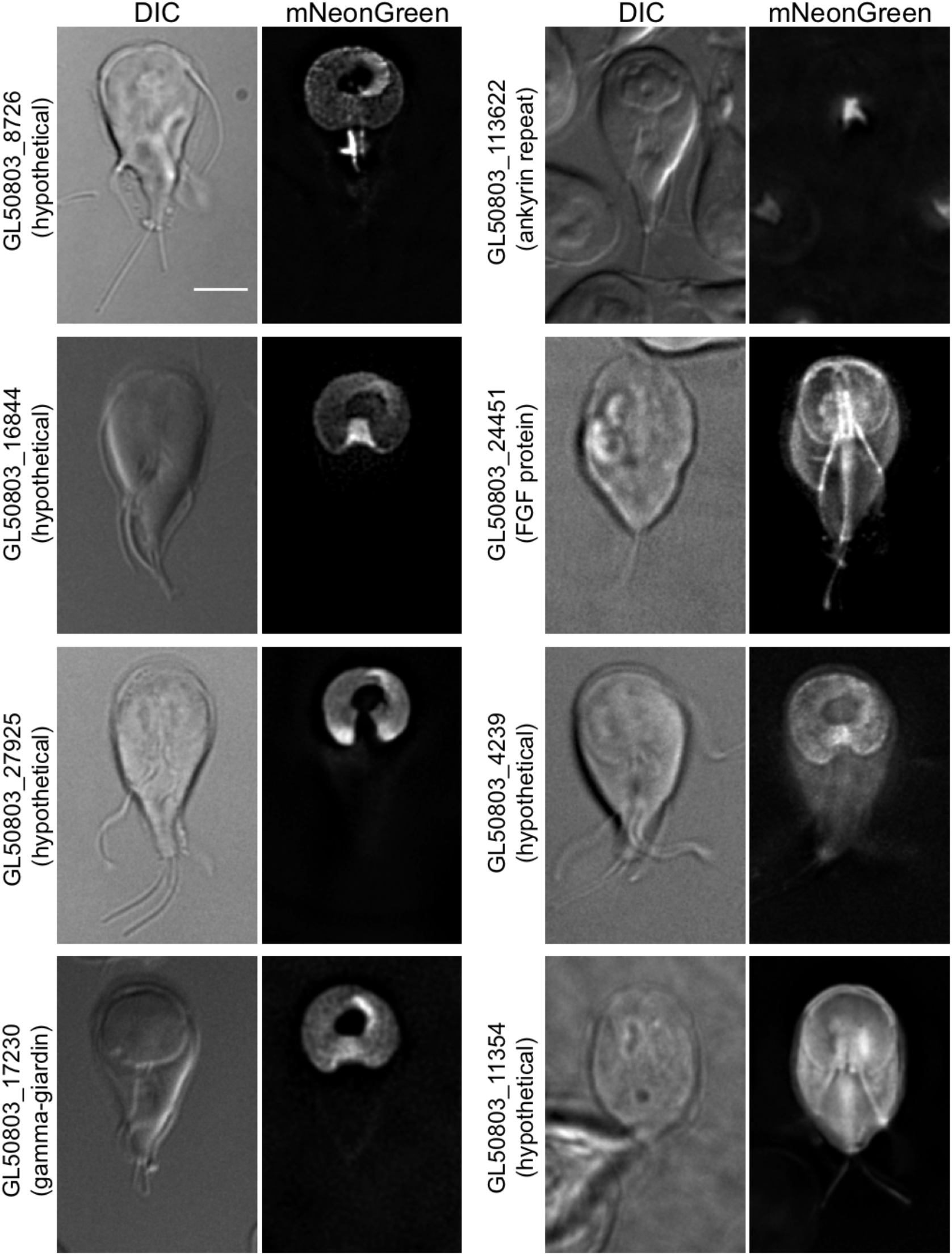
Putative *Gl*Actin Interactors with ventral disc localization. Eight proteins identified localized to the ventral disc based on mNeonGreen fusions. GL50803_8726 was more prominent on the disc margin and the overlap zone and appeared on the median body. GL50803_16844 was enriched in the ventral groove and the overlap zone; GL50803_1720 was similarly enriched in the overlap zone. GL50803_4239 had a slightly brighter signal in the ventral groove, the only place where GL50803_113622 localized. GL50803_11354 and GL50803_24451 faintly outlined both the ventral disc and the flagella. GL50803_27925 is excluded from the ventral groove. Scale bar, 5 μm.

### Co-localization of Candidate Interacts with *Gl*Actin

As there is no live *Gl*Actin marker, we performed immunofluorescence assays on one protein per category of localization with an anti-*Gl*Actin antibody to co-localize them (Figure 9). Phalloidin does not bind to *Gl*Actin (6), so polymerized *Gl*Actin is difficult to distinguish from pools of monomeric *Gl*Actin in fixed cells. Similarly, because *Gl*Actin is ubiquitously distributed throughout the cell, co-localization is not easily discerned. We used the JACoP plugin from ImageJ to evaluate co-localization. Based on Pearson’s Correlation Coefficients (PCC), the most prominent co-localization between *Gl*Actin and its putative interactors can be seen in the internal membrane protein GL50803_22855 and GL50803_13774, which localizes to the flagellar pores. The marginal plate protein GL50803_9030 and the nuclear localized GL50803_102813 also had PCCs > 0.5, indicating co-localization.

**Figure 9:**
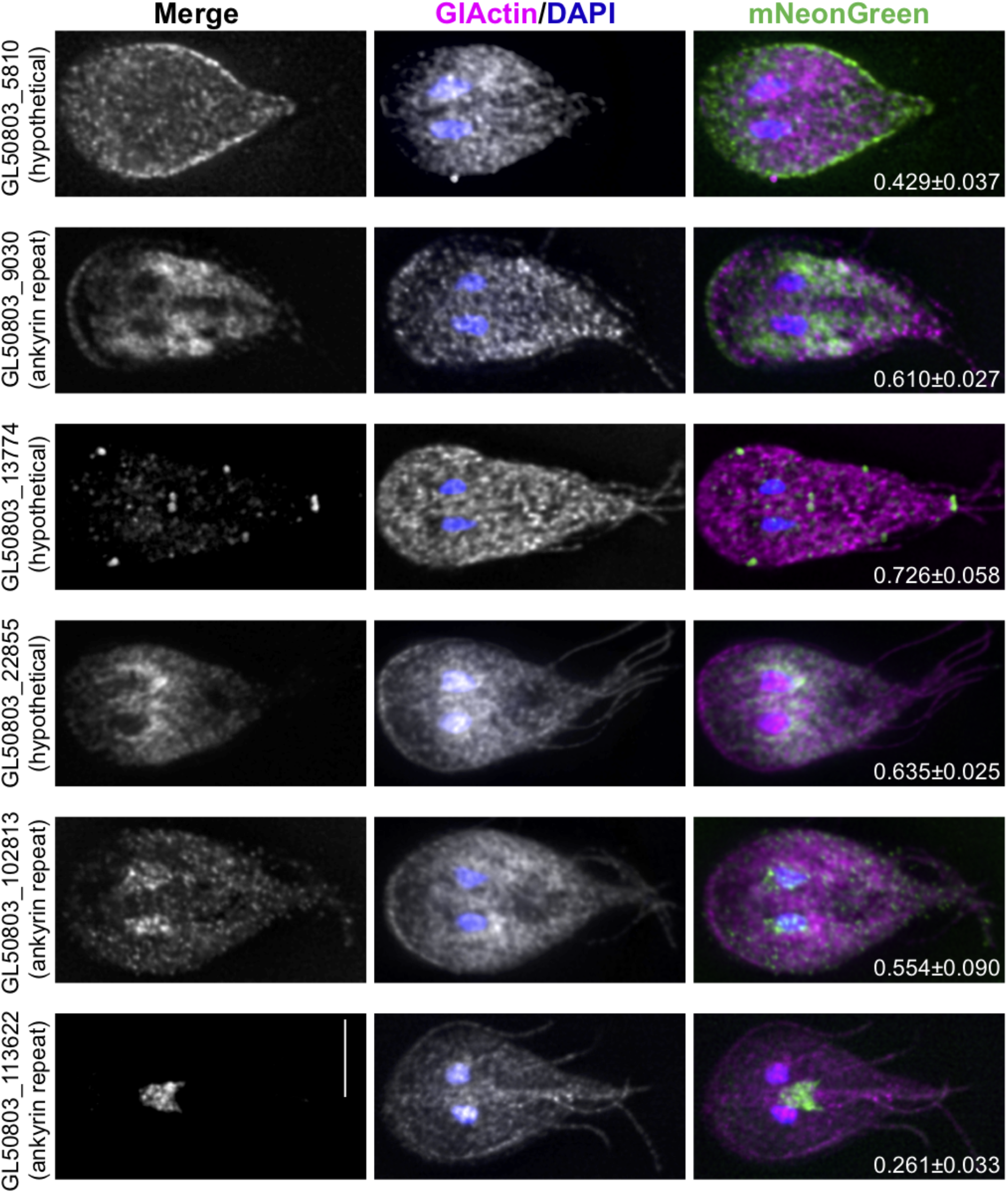
Immunostained *Gl*Actin is localized throughout the cell. One protein from each category of localization was co-localized with *Gl*Actin in fixed *Giardia* trophozoites. Nuclei are stained with DAPI (blue), *Gl*Actin is magenta, and each mNeonGreen-tagged interactor is in green. Average projections are shown. Scale bar, 5 μm. Pearson’s correlation coefficients using Costes randomization were calculated for three images of each protein, shown as mean±SD on each merge. The Costes P-value was 1 for all candidates.

In order to validate association with *Gl*Actin for a subset the identified proteins, we made 3xHA constructs to perform co-immunoprecipitations. We chose one protein per category based on its presence in all three replicates. The exception is GL50803_8854, as none of the proteins from the marginal plate category were found in all three replicates. We chose GL50803_8854 based on its high expression and presence at both the marginal plate and flagellar pores, structures where *Gl*Actin is enriched. We were able to confirm complex formation between *Gl*Actin and GL50803_27925 (disc), GL50803_8854 (marginal plate), and VPS29 (plasma membrane). Association between *Gl*Actin and GL50803_12999 (internal membranes) was ambiguous (Figure 10), possibly due to the latter’s association with membranes and/or low expression. The nuclear interactor GL50803_33989 did not express at high enough levels to be visible on a western blot (data not shown). Our co-immunoprecipitation results confirm that we have identified *Gl*Actin-associated proteins, increasing our confidence in the data presented here. It should be noted, however, that despite its co-localization with *Gl*Actin, the flagellar pore protein GL50803_13774 did not co-immunoprecipitate with *Gl*Actin. This does not necessary indicate that they do not interact; the 3xHA tag could have intefered with the interaction. Another possibility is that the interaction is too weak to be detected by western blot, since LC-MS/MS is a more sensitive technique.

**Figure 10:**
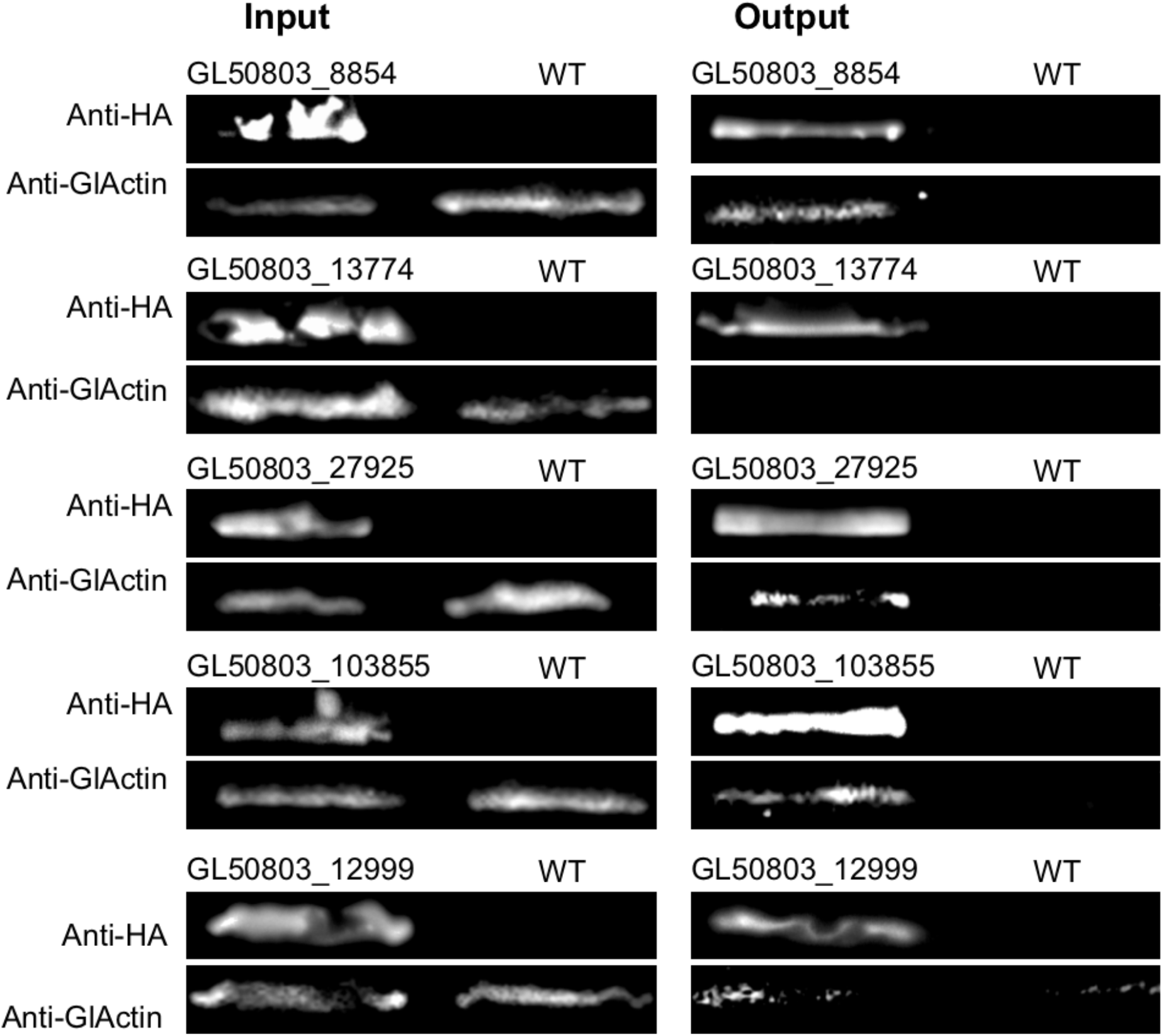
Reciprocal co-immunoprecipitation. One protein from each category of localization was tagged with 3xHA and immunoprecipitated, followed by anti-*Gl*Actin Western blotting. Three of the six proteins tagged definitively complexed with *Gl*Actin, including GL50803_8854 (marginal plate), Gl50803_103855 (plasma membrane), and GL50803_27925 (ventral disc).

## Discussion

We performed LC-MS/MS to identify putative *Gl*Actin interactors and found them to localize to multiple regions within the cell, including the nuclei, plasma membrane, endomembrane system, marginal plate, flagella, and ventral disc. We also found a number of previously validated *Gl*Actin-binding proteins (Supplemental Table 1) as well as several proteins which had already been localized in live cells by the Dawson lab (UC Davis) as part of the *Giardia* genome database project (Supplemental Table 2) (17). Seventeen of the proteins in this screen have no recognizable identity and lack conserved domains; thus, we were unable to find any clues to their function within their sequence. We also used I-TASSER to search for structural homologs, but this search too yielded no results (35). Six proteins identified in our screen contained ankyrin repeat domains, scaffolds for protein recognition and interaction, which are present in actin regulators in other organisms (37, 38). We also found four alpha-giardins, annexin-like cytoskeletal proteins unique to *Giardia* (39). The membrane-localized alpha-6 giardin may have a role in membrane organization, similar to the canonical annexin A-II, which binds F-actin and mediates its interactions with membranes (40, 41).

Nuclear actin has been described in other eukaryotes, including mammals, plants, insects, and other protozoa (42). Five of our putative F-actin interactors localized to *Giardia’s* two nuclei. When immunostained, *Gl*Actin is visible in the nuclei; our lab previously demonstrated roles for *Gl*Actin in nuclear positioning and size (6, 10, 13).

Both monomeric and filamentous actin have functions within the nucleus. G-actin has a role in gene regulation through formation of the pre-initiation complex and in chromatin remodeling through its association with the BAF complex (42, 43). In contrast, F-actin has a role in intranuclear mobility, WNT-dependent transcription, and nuclear stability (14, 42, 44). Notably, most of these functions depend on nuclear myosins, which are missing from the *Giardia* genome (3); however, it is still possible that *Gl*Actin fulfills these functions, particularly since a number of our putative interactors localize to the nuclei.

In model organisms, actin also works not only within but immediately around the nucleus for protection, as well as anchoring the nucleus and in nuclear positioning during mitosis, mediated by the Linker of Nucleoskeleton and Cytoskeleton (LINC) complex (44, 45). While *Giardia* lacks the LINC complex (6), it is likely that actin is still involved in nuclear positioning, since *Gl*Actin knockdowns results in mislocalized nuclei (6).

In model eukaryotes, the Arp2/3 complex drives actin polymerization to mediate clathrin-coated pit internalization, which is nucleated by the AP-2 complex (46). Although *Gl*Actin’s role in membrane trafficking has been demonstrated, its possible involvement in clathrin-mediated endocytosis remains controversial (47–49). However, the presence of the putative clathrin light chain among our interactors points to some involvement. Recent evidence also suggests *Gl*Actin interacts with clathrin heavy chain (50). Due to the absence of most conserved endocytic proteins, the mechanism for clathrin-mediated endocytosis in *Giardia* is poorly understood. The clathrin heavy chain has been demonstrated to interact with components of the AP-2 adaptor complex, specifically alpha and beta adaptins, which localize to the plasma membrane (51). The mu subunit of AP-2 has been localized to the hybrid endosome/lysosome peripheral vacuoles and plasma membrane (52). It was therefore unexpected to see sigma adaptin, central to the structure of the AP-2 complex (53–55), localize to the nucleus in addition to the plasma membrane. However, some of the other adaptins, notably alpha adaptin, have a function in nuclear translocation (56); it is possible that both sigma adaptin and *Gl*Actin are involved in this process as well, particularly since sigma adaptin has both nuclear and plasma membrane localization.

Two other membrane-localized proteins, vacuolar protein sorting (VPS) 29 and 35, are subunits of the cargo recognition particle of the retromer complex. Retromer is involved in the endosome-to-Golgi retrograde transport pathway and mediates the recycling of membrane receptors in yeast and mammals (57). Previous work in *Giardia*, which shows VPS29 localizing to the ER, conflicts with our results; however, the previous study used an HA-tagged protein that was localized in fixed cells. However, the same study localized VPS35 to the plasma membrane (58), so it is expected that at least a subset of VPS29 would also localize to the plasma membrane. While VPS26, the other subunit of the cargo recognition particle, did not fit our criteria for possible F-*Gl*Actin interactors, it did appear in our screen (Supplemental Table 3). Since there is no Golgi in *Giardia* and the sorting nexins which constitute the other retromer subunit are also missing from the *Giardia* genome, the role of *Giardia’s* cargo recognition particle is unknown. VPS35 has been shown to bind to hydrolase, but its function in doing so was not explored (58). Furthermore, while the canonical VPS35 is known to coordinate with actin through interaction with FAM21 and the WASH complex (59), these are also absent in *Giardia*.

While the *Giardia* genome lacks moesin, ezrin, radixin, or any ERM domain proteins, *Gl*Actin does localize to the cellular cortex, so one or more of the putative interactors also localized here could fulfill the role of a cortical crosslinker. Furthermore, *Gl*Actin-depleted cells have defects in cell polarity and cell shape (6). Another possible role for *Gl*Actin interactors localized to the plasma membrane is in phagocytosis, a process previously unrecorded in *Giardia*, but for which evidence has recently been published (60); if further research shows *Giardia* to be capable of phagocyotis, *Gl*Actin is likely to be involved (60, 61), given its role in phagocytosis in other organisms.

In addition to their presence in the plasma membrane, both the alpha-giardin GL50803_14551 and the hypothetical protein GL50803_17255 also extended into the ventrolateral flange (VLF), the hypothesized membrane reservoir that skirts *Giardia* cells. Short *Gl*Actin filaments appear in the VLF, and another novel actin interactor, Flangin, has been implicated in its function (32); it is likely that more of these interactors are important for maintenance and function of the VLF.

Many of our newly identified interactors localize to axonemes. Actin is a key component of the flagella inner dynein arm, although the actin within the inner arm is monomeric and complexed with a p28 homodimer (62); additionally, actin appears to play a structural role in γ-TURC (63). Immunostaining indicates *Gl*Actin to be within the flagella and we have previously identified p28 as an interactor (20, 64). Actin’s role in the flagella of other organisms is not merely structural; for instance, actin polymerization is required for flagellar biogenesis and intraflagellar transport in *Chlamydomonas reinhardtii* (65, 66). Morpholino knockdown of *Gl*Actin results in flagella which are mispositioned or even missing, implicating *Gl*Actin in flagellar positioning (6). The two flagellar pore interactors we identified could also be involved in this process; future studies could investigate the role of these proteins through depletion to test if resulting flagella lack stability, are trapped within the cell body, or exit the cell at an incorrect location.

Despite the reduction of *Giardia’s* endomembrane system, it has a relatively conventional ER, and two of our proteins had ER-like localizations, consistent with *Gl*Actin’s previously established role in membrane trafficking. Two others, GL50803_12999 (hypothetical) and GL50803_8044 (7 transmembrane domain) localize to an undescribed compartment. Since *Giardia* has unique organelles such as the Golgi-like encystation specific vesicles, it is likely that more compartments exist in *Giardia* than are currently recognized and these proteins may be involved in their function.

The marginal plate, a crescent-shaped structure at the anterior of the cell, has not yet been functionally studied, limiting our ability to infer the function of the proteins located there. While it has been suggested to be involved in attachment, this role is speculative. Since immunofluorescence assays show enrichment of *Gl*Actin at this site, it is possible that the meshwork of filaments seen in electron micrographs (67)are composed of *Gl*Actin or that *Gl*Actin acts upon this structure. Intriguingly, after depletion, *Gl*Actin localization persists at the marginal plate, suggesting that the associated *Gl*Actin is highly stabilized (13).

Of all the structures to which these putative *Gl*Actin interactors localize, the ventral disc is perhaps the most surprising. One previous study localized *Gl*Actin to the ventral disc (68); however, this study used a heterologous antibody, raised against chicken gizzard actin, for immunostaining. This finding is almost certainly an artifact, given that the same study found similar localization patters for actin-binding proteins not present in the *Giardia* proteome (68). While the disc itself is composed of microtubules, their polymerization dynamics are not responsible for flexation and/or attachment; it is possible that *Gl*Actin, as a force-generating protein, plays a role in this process. *Giardia’s* ability to attach, and thus its health, depends on the disc, and as this structure is unique to *Giardia*, many of the proteins that localize there have no relationship to proteins in other organisms. It is noteworthy that the different disc proteins had different localizations, indicating that the different regions of the disc may have different relationships to *Gl*Actin. Perhaps most notable are GL50803_16844 and GL50803_113644, as they are highly enriched in the margin near the overlap zone and/or the ventral groove; these regions are hypothesized to regulate fluid flow beneath the ventral disc (69). While *Gl*Actin does not appear concentrated in the ventral disc in immunofluorescence images, this could be due to steric hindrance between the antibody and proteins on the ventral disc, similar to the faint tubulin immunostaining seen in the disc compared to the amount of signal observed when using fluorescently-tagged tubulin (36). Another possibility is that F-actin is localized to the disc, but it is obscured by the pool of monomeric *Gl*Actin in our immunostaining. Defects in one of our disc-localized interactors, gamma-giardin, results in misshapen discs, but normal adherence, suggesting the role of gamma-giardin to be structural rather than actively mechanical (70). Notably, over half of our interactors with previously reported localizations are at the ventral disc (Supplemental Table 2) (17, 71).

Here we have characterized a number of putative *Gl*Actin interactors, with a diversity of localizations and possible applications. Study of *Gl*Actin is challenging as we still lack a live marker for actin in *Giardia* and markers that generally recognize actin in other eukaryotes, such as LifeAct, F-tractin, and Utrophin, do not label filamentous structures in *Giardia*. The proteins identified in our study could be truncated to potentially be used as a live *Gl*Actin marker, but development of such a tool would be difficult in those which lack identifiable domains. Of particular interest are the proteins which localize to the ventral disc, the ventrolateral flange, and the marginal plate. If, as hypothesized, the plate is important for attachment, this work has potentially uncovered a connection between *Gl*Actin and all of *Giardia’s* mechanisms for attachment, which is required for maintaining infection. Thus, there are ample opportunities for drug development in further study of these proteins, particularly since few of them have homologues in humans.

## Materials and Methods

### Parasite Strain and Growth Conditions

*G. lamblia* strain WB clone 6 (ATCC 50803; American Type Culture Collection) was cultured under standard conditions (72).

### Lysis and Crosslinking

3 L of wild type *Giardia* trophozoite culture was grown to confluence in 1 L bottles as described previously (10). Cells were iced for 2 hours to induce detachment, then centrifugation for 30 minutes at 1500xg at 4° C, then washed twice with 1XHBS (HEPES buffered saline) plus 1xHALT protease inhibitor cocktail (Thermo Scientific Catalog number 78430) by resuspension and centrifugation for 15 minutes at 1500xg at 4°C. Each culture was then resuspended in 1.6 mL of buffer. Two were resuspended in F-buffer (20 mM HEPES, 0.2 mM CaCl2, 80 mM KCl, 10 mM imidazole, 1 mM MgCl2, 1 mM EGTA, 5% Glycerol, 10 mM ATP, 1X HALT protease inhibitors, pH 7.2) while one was resuspended in G-Buffer (20 mM HEPES, 0.2 mM CaCl2, 0.2 mM ATP, 1X HALT protease inhibitors, pH 7.2).

One of the F-buffer cultures was then treated with crosslinker DSP (dithiobis(succinimidyl propionate)) (Thermo Scientific Catalog number 22585) to a final concentration of 1 mM and incubated for 30 minutes at room temperature, quenched with Tris-HCl pH 7.5 to a final concentration of 20mM and incubated at 15 minutes, then lysed in the manner described below. The other F-buffer culture and the G-buffer culture were lysed with sonication, pulse for 25 seconds each at 20% power with 1-minute rest between each sonication, repeated a total of 4 times minimum. Sonicated cells were allowed to rest on ice for 30 minutes, then cleared with a 10,000xg spin for 10 minutes at 4°C. The supernatant of these two cultures was then treated with DSP crosslinker to a final concentration of 1 mM and incubated for 30 minutes at room temperature, quenched with Tris-HCl pH 7.5 to a final concentration of 20mM and incubated at 15 minutes. This was repeated for a total of three independent replicates per condition.

### Purification and Sample Preparation for Mass Spectrometry

Lysates were incubated overnight with an anti-*Gl*Actin antibody (6) at 4°C, then incubated with 500 μL Pierce™ Protein A agarose (Catalog number 20333) bead slurry and incubated with end-to-end mixing for 2 hours at room temperature. A total of four washes were performed by addition of 0.5 mL of wash buffer (G- or F-Buffer with an additional 150 mM NaCl), vortexed, and centrifuged for 2-3 minutes at 2500xg. For elution, 250 μL of 0.2 M Glycine pH 2.5 was added to the beads and incubated for 5 minutes, centrifuged for 3 minutes at 2500xg, and the supernatant was collected. This step was then repeated and the two eluate fractions were combined. The pH was then neutralized through addition of 50 μL 1 M Tris-HCl pH 7.5.

For each condition, 5X sample buffer (0.225 M Tris-HCl, pH 6.8, 50% glycerol, 5% SDS, 0.05% bromophenol blue, 0.25 M DTT) was added to a final concentration of 1X subsequent to running on a 10% SDS-PAGE gel and stained with SYPRO Ruby (Pierce). Bands corresponding to the heavy and light antibody chains were excised from the gel and discarded; the remainder of the gels were divided into rectangles of approximately equal size. These rectangles were cut into squares of approximately 1-2 mm, washed with 200 μL water, then incubated with 200 μL of 50% acetonitrile and 25 mM ammonium bicarbonante for 5 minutes. After addition of another 200 μL acetonitrile and further incubation for 1 minute, the gel was dried with a speed vacuum, followed by another addition of 50 μL of 25 mM ammonium bicarbonate and 10 mM TCEP. Samples were then incubated at 60°C for 1 hour, after which the supernatant was removed and replaced with 50 μL of 25 mM ammonium bicarbonate and 10mM iodacetamide, then incubated for 20 minutes in the dark. Another wash was performed with 400 μL water and the 5-minute incubation with 200 μl 50% acetonitrile, 25 mM ammonium bicarbonate and following steps were repeated. Protein samples were then digested with 20 μL of 0.01 μg/μL Promega Trypsin in 25 mM ammonium bicarbonate. More ammonium bicarbonate was added to cover, followed by overnight digestion. Supernatant was then removed for LC/MS-MS analysis. Protein was further extracted from the gels by addition of 50 μL of acetonitrile, vortexing, and removing and saving supernatant. One last extraction was performed by addition of 50 μL 60% acetonitrile, 0.1% formic acid, vortexing for 5 minutes, and then removing the supernatant. The supernatants were then combined, dried, and resuspended in solvent A, then analyzed by LC/MS-MS as described below.

### Liquid Chromatography and Mass Spectrometry

Liquid chromatography – mass spectrometry was performed on a Velos Pro (Thermo) with an EasyLC 1000 HPLC and autosampler (Thermo). Samples were solubilized in loading buffer (0.1% trifluoroacetic acid and 2% acetonitrile in water), and 6 ul was injected via the autosampler onto a 150-μm Kasil fritted trap packed with Reprosil-Pur C18-AQ (3-μm bead diameter, Dr. Maisch) to a bed length of 2 cm at a flow rate of 2 ul/min. After loading and desalting using a total volume of 8 ul of loading buffer, the trap was brought on-line with a pulled fused-silica capillary tip (75-μm i.d.) packed to a length of 25 cm with the same Dr. Maisch beads. Peptides were eluted off the column using a gradient of 2-35% acetonitrile in 0.1% formic acid over 90 minutes, followed by 35-60% acetonitrile over 5 minutes at a flow rate of 250 nl/min. The mass spectrometer was operated using electrospray ionization (2 kV) with the heated transfer tube at 200 C using data dependent acquisition (DDA), whereby a mass spectrum (m/z 400-1600, normal scan rate) was acquired with up to 15 MS/MS spectra (rapid scan rate) of the most intense precursors found in the MS1 scan.

Database searches were performed using Comet (73) searched against the protein sequence database GiardiaDB 3.1_GintestinalisAssemblageA_AnnotatedProteins.fasta to which was appended sequences of common contaminants (e.g., human keratins). The peptide mass tolerance was 3 Da, and the fragment ions were sorted into 1 Da bins (roughly corresponding to a fragment ion mass tolerance of +/- 0.5 Da). Semi-tryptic cleavages and up to two missed tryptic cleavage sites were allowed. Oxidized methionine and acetylation of the protein N-termini were allowed as variable modifications. Carbamidomethylated cysteine was a fixed modification. False discovery rates (1%) were determined using PeptideProphet (74), and protein inferences were made using ProteinProphet (75). Spectral count analysis was performed using Abacus (76).

### Data Availability

The mass spectrometry proteomics data have been deposited to the ProteomeXchange (77) Consortium via the PRIDE partner repository with the dataset identifier PXD026067 (78). Reviewer account details: Username: Password: v05IifYI

### Bioinformatics

Delta-BLAST, Protein BLAST, and Pfam searches were used to identify homologues and conserved domains for each putative interactor. I-TASSER was used to search for structure-based function prediction.

### Vector Construction

All constructs in this study were C-terminal fusions with the exception of the N-terminally tagged GL50803_33989 and GL50803_7323, using Gibson reactions with linearized vectors and PCR products (7, 79). For primer sequences and workflow, see Supplemental Table 4.

### Immunofluorescence Microscopy

Fixation and imaging were performed as described in previous work (80). FIJI/ImageJ were used for image analysis, including co-localization analysis by Just Another Colocalization Plugin (JACoP) (81).

### Live Cell Imaging

Cells were chilled with ice for 15 minutes to detach from the culture tube and then placed into an Attofluor cell chamber (Molecular Probes) and incubated in a GasPak EZ anaerobic pouch (BD) or a Tri-gas incubator (Panasonic) set to 2.5% O2, 5% Co2 for 90 minutes at 37° C. Cells were then washed four times with HEPES-buffered saline (137mM NaCl, 5mM KCl, 0.91mM Na2HpO4-heptahydrate, 5. 55mM Glucose, 20mM HEPES, and pH7), overlaid with a mixture of 0.7% ultra-low gelling agarose (Sigma A2576) melted in HEPES-buffered saline, cooled for 10 minutes at room temperature to solidify, then imaged. Live cell imaging was performed on a DeltaVision Elite microscope (GE) equipped with DIC optics, using a 100X 1.4 NA or 60X 1.42 NA objective, and a sCMOS 5.4 PCle air-cooled camera (PCO-TECH).

### Co-immunoprecipitation

500 mL of *Giardia* cell cultures were grown for 3 days, then iced for 2 hours to detach and spun for 1500xg at 4 °C. Cells were then washed twice in HBS with 2XHALT protease inhibitors and 10 μM chymostatin, 1μM leupeptin, and 1 μM E64. Each pellet was resuspended to a final volume of 1.2 mL and 100 mM DSP in DMSO was added to a final concentration of 1 mM and incubated at room temperature for 30 minutes. The reaction was quenched for 15 minutes with an addition of Tris-HCl pH 7.4 final concentration 20 mM. Cells were then pelleted by spinning for 7 minutes at 700xg and resuspended in 350 μL lysis buffer (80 mM KCl, 10 mM imidazole, 1 mM MgCl2, 1 mM EGTA, 5% Glycerol, 20 mM HEPES, 0.2 mM, CaCl2, 10 mM ATP, 0.1% Triton X-100, 500 mM NaCl, pH 7.2).

Cells were then lysed by sonication as and cleared as described above. A volume of 17.5 μL of equilibrated EZview Red Anti-HA Affinity gel (Sigma) was added to each tube of lysate, then incubated at 4°C with end-over-end mixing for 1 hour. Beads were then spun at 8,200xg for 30 seconds and the supernatant was discarded, followed by a total of three washes with 750 μL wash buffer (80 mM KCl, 10 mM imidazole, 1 mM MgCl2, 1 mM EGTA, 5% Glycerol, 20 mM HEPES, 0.2 mM CaCl2, 10 mM ATP, 0.5% Tween, 500 mM NaCl, pH 7.2). Each wash consisted of end-over-end rotation for 5 minutes followed by a 30 second spin at 8,200xg. Protein was then incubated with 50 μL of 8 M Urea at RT for 20 minutes to elute, followed by addition of 5X sample buffer (QIAGEN) to a final concentration of 1X, boiled for 5 mins at 98° C, and run on 12% SDS-PAGE gel, followed by Western blot protocol described previously (7).

---

**Supplemental Table 1: Previously validated *Gl*Actin interactors identified in this screen**

**Supplemental Table 2: F-*Gl*Actin interactors which have been previously localized**

**Supplemental Table 3: Complete list of proteins identified in this screen as *Gl*Actin interactors**

**Supplemental Table 4: Primer sequences and workflow for tagging each protein**

## Acknowledgements

We thank Kelly Hennessey, Kelli Hvorecny, Han-wei Shih, Elizabeth Thomas, and Germain Alas for manuscript editing.

This material is based upon work supported by the National Science Foundation Graduate Research Fellowship under Grant No. DGE-1762114.

